# An elective course for medical students on innovation and entrepreneurship

**DOI:** 10.1101/148569

**Authors:** Erik Reinertsen, Arun Mohan, Angela Fusaro

## Abstract

Here we describe “Innovation and Entrepreneurship in Medicine” (IEMed), a short elective course that exposed medical students to innovation and entrepreneurship. We assess students’ self-rated familiarity with various learning objectives after the course and report survey outcomes. Students were most interested in further educational offerings focused on startups and innovation. However, students self-reported relatively lower levels of understanding in regulatory, reimbursement, and legal aspects of health tech. Most participants reported a desire to collaborate with or consult for industry partners, rather than to found or work at a startup. We conclude with a broader discussion of the need to expose medical students to opportunities in the technology industry.

## 1 Introduction

Entrepreneurial and software-driven technology companies, known as “tech”, are reinventing established industries, including transportation, national defense, and even healthcare. For the past three years, investment in digital health – a broad category of tech including analytics, biosensing, population health management, medical records, and software-driven devices – has exceeded $4B annually [1]. However, implementation of technology into clinical workflow has been slow.

Medical students are positioned to contribute to tech’s foray into biomedicine for two reasons. First, they are comfortable with technology, which permeates most aspects of their lives as consumers. Second, they possess insight about unmet clinical needs and care delivery. Unfortunately, medical students are not exposed to the tech industry even though efforts therein can represent “translational research in its rawest form” [2].

## 2 Approach

To address this problem, we created an elective course called “Innovation and Entrepreneurship in Medicine” (IEMed) at our institution - a private medical school in a large metropolitan city in the United States. This course was for second-year medical students, held in the fall semester, and lasted three months. We developed the following learning objectives:

1. Assess a startup using the business model canvas as a framework.
2. Learn about healthcare economics and reimbursement.
3. Appreciate how behavioral economics assesses decision-making.
4. Review advances in digital health and health information technology.
5. Survey accelerators, incubators, and other startup resources.
6. Learn about patents and the fundamentals of intellectual property.
7. Understand the role of angel, seed, and venture capital investors.
8. Meet like-minded students, clinicians, engineers, and startup founders.
9. See how academic clinicians collaborate with the tech industry.

Nine students enrolled in 2014, and ten students enrolled in 2015. Each learning objective was explored through a lecture and Q&A featuring a guest speaker with relevant experience. Sessions were also accompanied by suggested readings. We recorded videos of several sessions to provide educational assets to a broader audience. The full course schedule, reading materials, and videos of select sessions were available through a course website at http://erikreinertsen.com/iemed.

Students also completed a project in teams of three. Each team selected and evaluated a startup company using the “business model canvas”: a visual chart describing market strategy, value proposition, infrastructure, customers, finances, etc. (Figure 1) [3]. Each week, teams focused on one component of the canvas, learned more about their startup by conducting research online, or speaking with founders of the startup and/or potential customers, and shared their findings with the class. We also connected medical students with MBA students at our institution’s business school to provide input. At the end of the course, teams presented their canvases and discussed clinical implementation and business development strategies.

**Figure 1:**
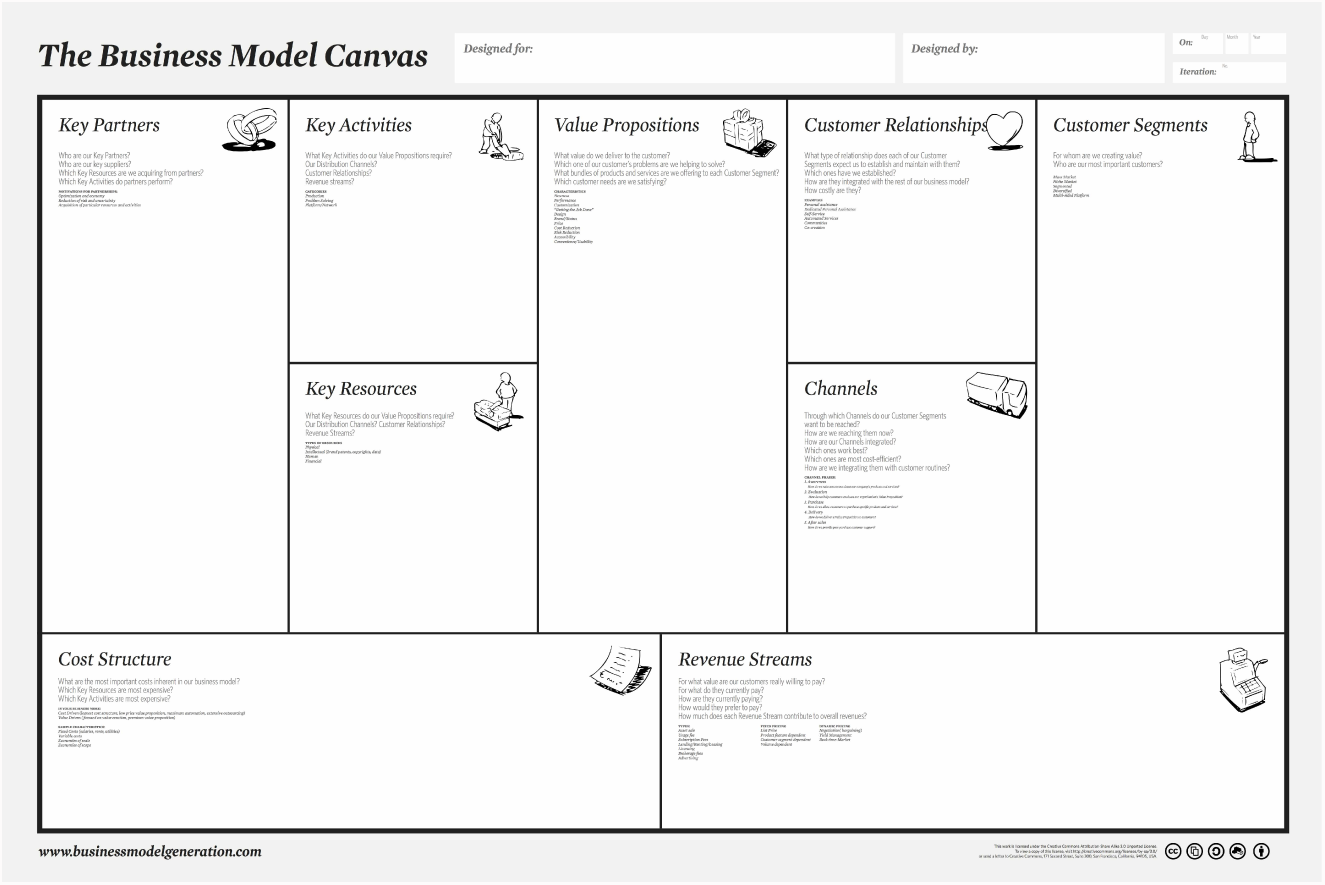
The business model canvas serves as an overview of major components of a startup company.

## 3 Outcomes

In the second year of IEMed, we administered a survey to enrolled students immediately after the last session to obtain feedback for subsequent course refinement (Figure 2). This study was exempt by our Institutional Review Board (IRB) as it fell under the category of “research on the effectiveness of or the comparison among instructional techniques, curricula, or classroom management methods”.

**Figure 2:**
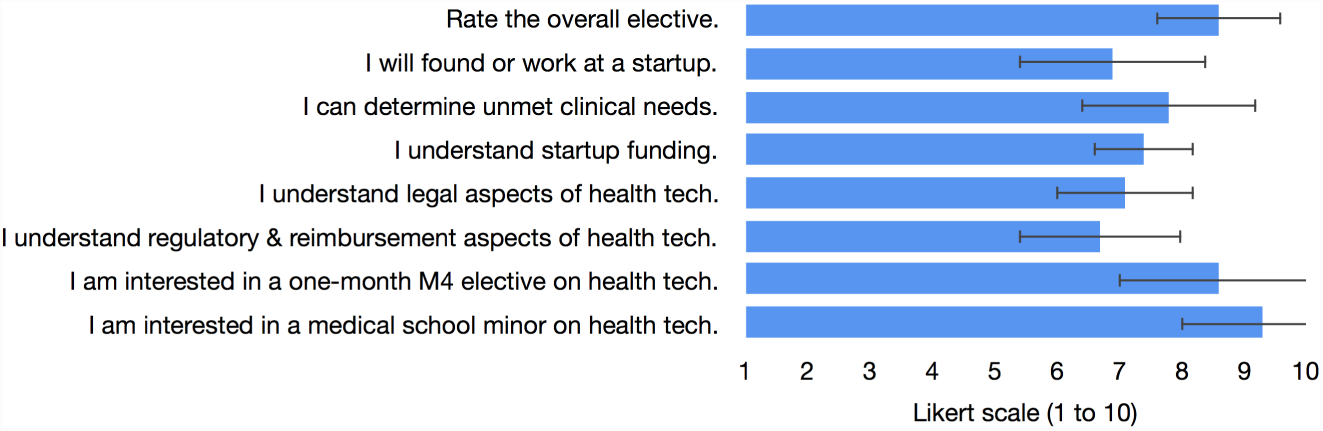
Results of survey administered at the end of IEMed investigating student perception of learning objectives, career interests, and future related educational offerings. Bar represents shows the mean, and the error bar shows the standard deviation.

Responders were asked to rate agreement with a statement using a Likert scale of 1 to 10, where 1 indicated strong disagreement, 5 indicated neutrality, and 10 indicated strong agreement. Ten students (100%) completed the survey, and 100% of students recommended the elective for future medical students. The three *highest* rated statements are shown in Table 1, and the three *lowest* rated statements are shown in Table 2.

**Table 1:** Survey results: highest rated statements

| Rank | Question | Mean ± std. deviation |
| --- | --- | --- |
| 1 | I am interested in a minor in startups and innovation | 9.3 ± 1.3 |
| 2 | I am interested in a one-month M4 elective on startups | 8.6 ± 1.6) |
| 3 | Rate the overall elective | 8.6 ± 1.0 |

**Table 2:** Survey results: lowest rated statements

| Rank | Question | Mean ± std. deviation |
| --- | --- | --- |
| 1 | I understand regulatory and reimbursement aspects of health tech | 6.7 ± 1.3 |
| 2 | I will found or work at a startup | 6.9 ± 1.5 |
| 3 | I understand legal aspects of health tech | 7.1 ± 1.1 |

When asked about career goals, students reported the desire to pursue residency, practice clinically, and work with startups in advisory, consulting, or collaborative roles. Students also suggested ways to improve the elective, including networking events with business, engineering, and law students, a more detailed case study of a startup from idea to market, and lectures on implementing technology in the clinic. The business model canvas project was the least favored aspect of the course; several students requested more in-depth guidance on using the framework.

## 4 Discussion

Our survey faced several limitations. It was only designed to gauge interest in specific topics within our elective. We measured perceptions after the course and did not survey a control group. Furthermore, our sample size was small and the survey results were self-reported. Aware of these constraints, we simply utilized survey feedback to inform future medical education initiatives.

We believe academia can do more to support medical student and physician involvement in the creation and implementation of technologies that improve patient care. Relevant topics in business, law, and engineering should be available at least as an option in the curriculum without increasing the time required to complete medical school. Students should be encouraged to consult for, collaborate with, or intern at companies, especially early-stage technology startups.

Unfortunately, medical students are tacitly discouraged from pursuing endeavors outside of clinical work, basic science, or clinical research. Changing this perception remains a major cultural challenge, yet also presents an opportunity for academic medicine to define how technology advances the delivery of healthcare. We believe medical schools should support and facilitate the involvement of medical students in these translational efforts.

A few progressive academic institutions are breaking with tradition by supporting such educational and training opportunities at the intersection of healthcare, business, and technology. Medical students at Stanford University can become involved with StartX Med, which is “focused on accelerating the development of Stanford’s top medical entrepreneurs through experiential education and collective intelligence” [4]. Sling Health is a national incubator for biomedical technology commercialization; it started as a nonprofit organization sat Washington University St. Louis and recently expanded to several other cities as well as our institution [5]. The Icahn School of Medicine at Mt. Sinai offers internships to medical students with McKinsey and Company, Verily Life Sciences, and Deloitte Consulting [6].

To respond to student demand at our institution for further educational opportunities related to technology and innovation, we are piloting a new project-driven elective for fourth-year medical students. Over one month, students identify unmet healthcare needs by speaking with patients, providers, payers, and other stakeholders. Students also learn from discussions with experts in engineering, information technology, product development, and business. At the end of the elective, students propose a product or process addressing the unmet need. This approach has utility both as a didactic tool and an ideation method for quality improvement projects. Creative problem-solving skills are important for clinical care, research, and more directly translational work, yet underrepresented in current medical curricula.

## 5 Conclusion

In summary, the IEMed elective exposed medical students to topics at the intersection of technology and healthcare, which are underrepresented in medical curricula. The elective was well-received by all participants. IEMed required only a few hours per week of faculty and/or teaching assistant time, as it drew heavily on guest speakers with relevant expertise. This model is simple, could be easily implemented at other institutions, and may contribute to the professional development of future leaders in healthcare technology and delivery.

## Acknowledgements

The authors thank the administrative leadership of Emory University School of Medicine for the institutional and logistical support that helped make this elective possible.

